# Enrichment of genomic variation in pathways linked to autism

**DOI:** 10.1101/2020.10.19.346072

**Authors:** Francisco J. Esteban, Peter J. Tonellato, Dennis P. Wall

## Abstract

The genetic heterogeneity of autism has stymied the search for causes and cures. Even whole-genomic studies on large numbers of families have yielded results of relatively little impact. In the present work, we analyze two genomic databases using a novel strategy that takes prior knowledge of genetic relationships into account and that was designed to boost signal important to our understanding of the molecular basis of autism. Our strategy was designed to identify significant genomic variation within *a priori* defined biological concepts and improves signal detection while lessening the severity of multiple test correction seen in standard analysis of genome-wide association data. Upon application of our approach using 3,244 biological concepts, we detected genomic variation in 68 biological concepts with significant association to autism in comparison to family-based controls. These concepts clustered naturally into a total of 19 classes, principally including cell adhesion, cancer, and immune response. The top-ranking concepts contained high percentages of genes already suspected to play roles in autism or in a related neurological disorder. In addition, many of the sets associated with autism at the DNA level also proved to be predictive of changes in gene expression within a separate population of autistic cases, suggesting that the signature of genomic variation may also be detectable in blood-based transcriptional profiles. This robust cross-validation with gene expression data from individuals with autism coupled with the enrichment within autism-related neurological disorders supported the possibility that the mutations play important roles in the onset of autism and should be given priority for further study. In sum, our work provides new leads into the genetic underpinnings of autism and highlights the importance of reanalysis of genomic studies of complex disease using prior knowledge of genetic organization.

**Author Summary:** The genetic heterogeneity of autism has stymied the search for causes and cures. Even whole-genomic studies on large numbers of families have yielded results of relatively little impact. In the present work, we reanalyze two of the most influential whole-genomic studies using a novel strategy that takes prior knowledge of genetic relationships into account in an effort to boost signal important to our understanding of the molecular structure of autism. Our approach demonstrates that these genome wide association studies contain more information relevant to autism than previously realized. We detected 68 highly significant collections of mutations that map to genes with measurable and significant changes in gene expression in autistic individuals, and that have been implicated in other neurological disorders believed to be closely related, and genetically linked, to autism. Our work provides leads into the genetic underpinnings of autism and highlights the importance of reanalysis of genomic studies of disease using prior knowledge of genetic organization.

## Introduction

Autism is a complex neurological disorder characterized by restricted communication, impaired social interaction, and repetitive behavior. Diagnosis typically occurs by 4.3 years of age, with a current CDC estimate of approximately 1 in 59 children meeting criteria for autism spectrum disorder (ASD).

Heritability estimates of ~90% [1] for ASD imply a strong genetic component, making it one of the most highly heritable genetic disorders. Despite this high heritability, genetic association studies have been unable to detect variants explaining more than a negligible fraction of the phenotypic variability present in study populations.

Published genome-scale attempts to identify variants with significant association to ASD have been met with limited success. Weiss et al. [2] were not able to identify any individual single-nucleotide polymorphism (SNP) meeting genome-wide significance, but were able to detect a highly penetrant microdeletion/microduplication in the 16p11.2 region, present in approximately 1% of the cases studied. Examining many of the same families using an alternative genotyping platform, Wang et al. [3] uncovered a common variant in the region between two cell adhesion genes, CDH9 and CHD10, that met criteria for genome-wide significance after aggregating results from multiple studies. Despite these successes, our picture of autism-associated variation remains limited.

The inability to find signal associated with the genetic causes of autism may be partially attributable to the phenotypic heterogeneity of ASDs, as the behavioral variation is likely to be rooted in high genetic heterogeneity. It may also be due to technological bias as the genotyping platforms used were designed to detect common, rather than rare, variants. Additionally, purely data-driven approaches to the analysis of genotyping data tend to suffer from low power due to inadequate sample sizes and the severity of multiple test correction. This can be more pronounced in complex disorders like autism where it is likely that many variants of weak to moderate effect, which require exponentially larger sample sizes to detect, contribute to the onset and progression of the disorder.

The success of GWAS has been debatable [4–7], with studies suffering from insufficient samples sizes leading to low statistical power, significant findings in gene deserts which provide little, if any, biological insight, and minimal reproducibility of significant findings. As a consequence, the field has been motivated to develop alternative approaches to GWAS analysis that can circumvent some of the signal detection problems associated with purely data-driven approaches while elucidating biologically meaningful variation. Borrowing from the field of microarray data analysis, which suffers similar problems of multi-dimensionality and low signal-to-noise ratio, methods classifiable as knowledge-driven strategies have emerged to improve detection of real biological signal among previously elusive GWAS data [3, 8–15]. Several of these approaches have enjoyed considerable success. The strategy employed by the majority of these methods begins by assigning a significance level to individual genes based on the p-values computed for the GWAS data. Once significance has been determined at the gene level, approaches assess pathway significance using a variety of approaches, including ranking genes and testing for overrepresentation in the top of the ordered list, building protein-protein interaction (PPI) networks, and testing the distribution of genes within a pathway that contain nominally significant SNPs. However, some strategies use the SNP data directly and look at distributions of nominally significant SNPs within pathways or annotation classes. In many cases these approaches are helping to filter the search space of genetic variants associated with various diseases and, in general, strategies like these are helping GWAS achieve some of the potential of their original promise.

In the present study, we reanalyzed data from two GWAS of autism families with a similar approach designed specifically to handle family-based data and to detect previously masked signal among *a priori* defined biological concepts. By doing so, we were able to identify SNPs significantly associated with autism at a whole-genomic scale that had been previously missed by standard analytical methods. These SNPs mapped to genes that were highly enriched for neurological disease candidates and that exhibited significant differential expression in blood profiles of an entirely different population of autistic individuals. Our results reveal significant genotypic variation in an autistic population that is likely to be linked to the molecular pathology of the disorder, and also highlight the important of using knowledge-driven approaches to study GWAS data that had previously yielded limited signal.

## Results

We used two published genome-wide association studies (GWAS), Broad and CHOP, for our analyses (Table 1). From these, we acquired a total of 500,000 and 550,000 SNPs for 751 (2,883) and 943 (4,444) families, respectively (Table 2). The overlap in genomic coverage between the Broad and CHOP studies was low with less than 68,000 SNPs in common between the two platforms, thus the union provided significantly higher resolution than either study independently. After calculating nominal p-values using the family-based association test (FBAT) [16], we mapped all SNPs from both the Broad and CHOP studies to the MSigDB C2 and C5 gene set collections. This mapping resulted in 3344 SNP sets that ranged in size from 1-57,525 SNPs (mean 1,315.8), representing anywhere from 1-1,873 (mean 59.3) genes (Table 2). We then used these sets to search for significant differences in genotype frequencies associated with the population of autistic cases in our two samples.

**Table 1.** Summary of SNP genotypes used for analyses. Individual samples genotyped at both facilities were acquired from the Autism Genetic Resource Exchange (AGRE), resulting in a large overlap in individuals genotyped in both cohorts. However, the use of different genotyping platforms at each facility produced low overlap in SNP coverage between the two cohorts.

| **Cohort** | **Platform** | **No. of Pedigrees** | **No. of Individuals** | **No. of SNPs** |
| --- | --- | --- | --- | --- |
| Broad | Affymetrix 5.0 | 557 | 2,137 | 269,214 |
| CHOP | Illumina<br>HumanHap<br>550 | 727 | 3,322 | 264,543 |
| Overlap | — | 554 | 2,093 | 31,217 |

**Table 2.**
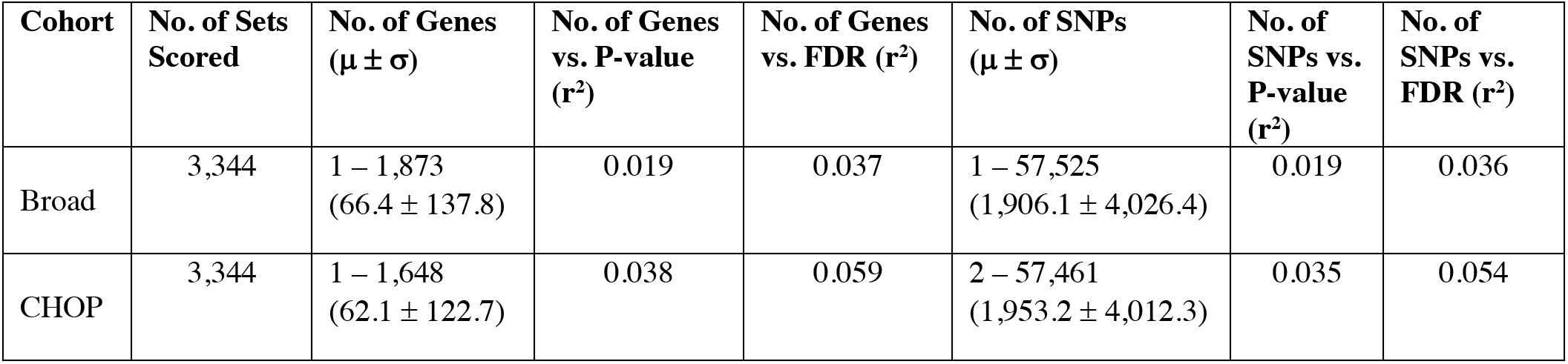
Summary of sets analyzed. The Broad Institute’s MSigDB C2 (curated) and C5 (GO) gene sets were used to group SNPs to enable knowledge-driven analyses of SNP genotypes. The resulting SNP sets ranged in size from 1 to 57,525 SNPs, representative of 1 to 1,873 genes. After performing analyses, Pearson correlation coefficients were calculated to ensure that set sizes did not bias the p-values; the strongest correlation found was not significant (r^2^ = 0.03).

### Set-based analysis

After excluding 7 SNP sets with low expected values and controlling the FDR at *q* = 0.05, 68 of the 3,344 (3.0%) sets tested achieved significance in either the Broad or CHOP data set (Table 3; Supplementary Table 1).

**Table 3.**
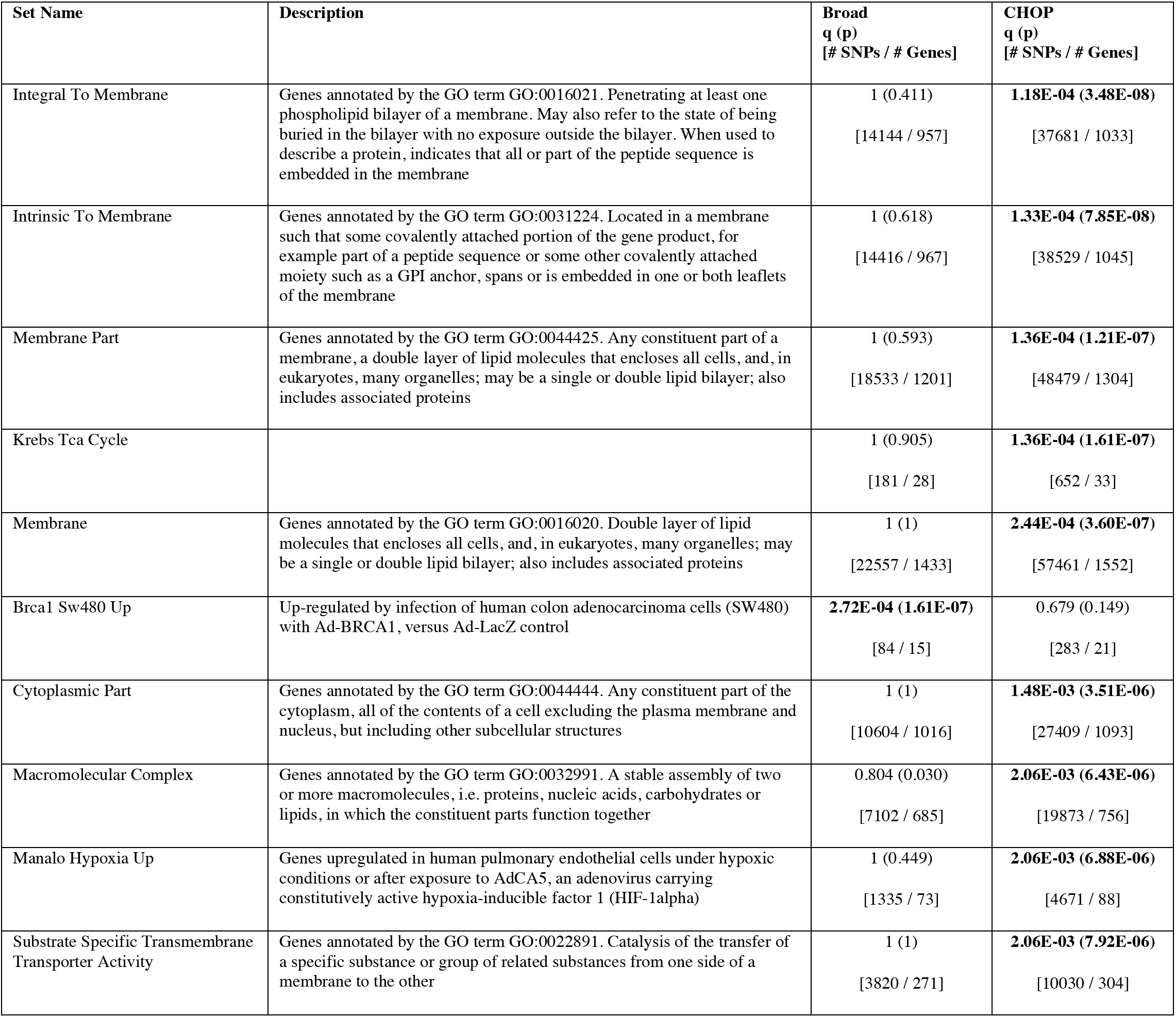
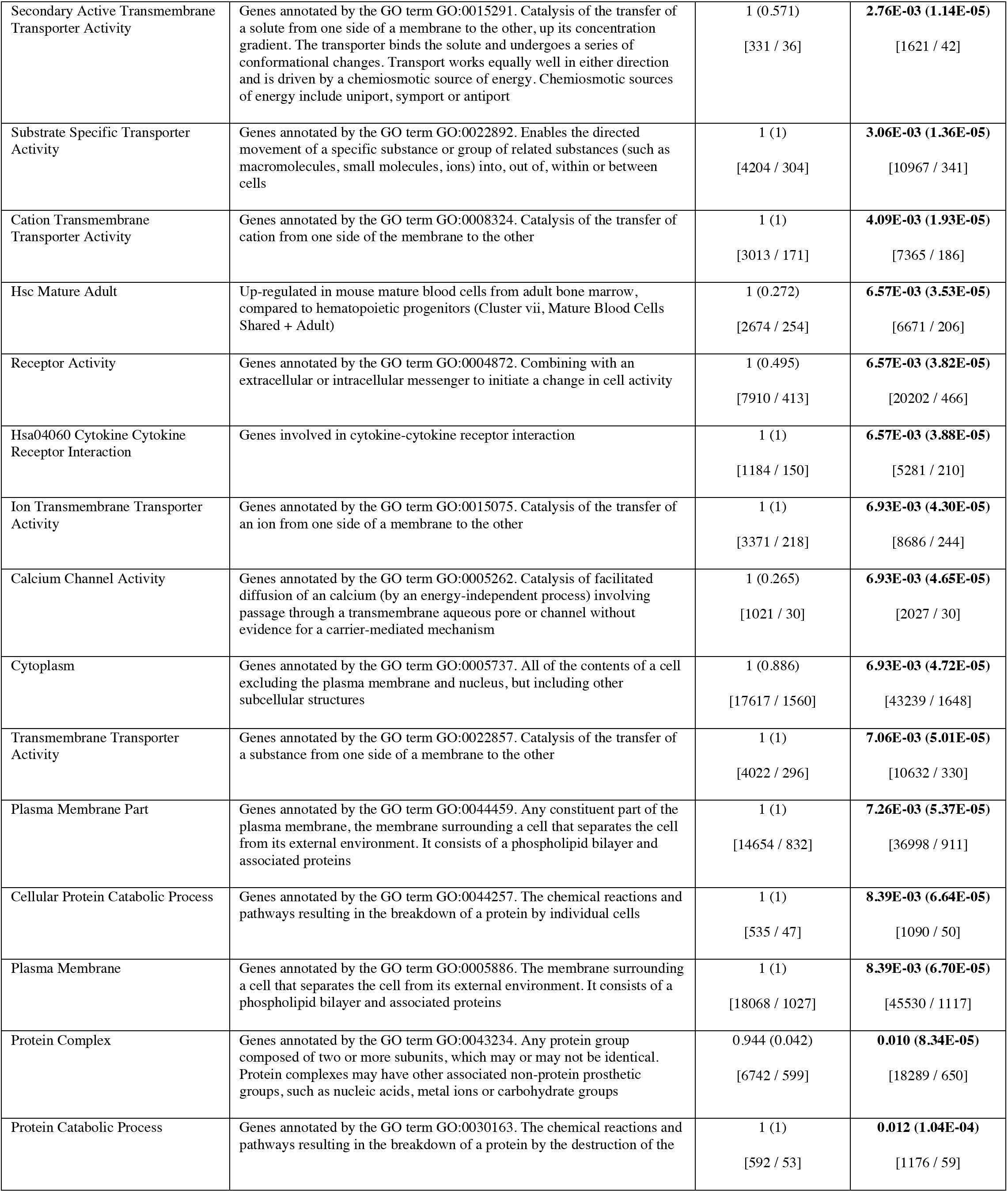
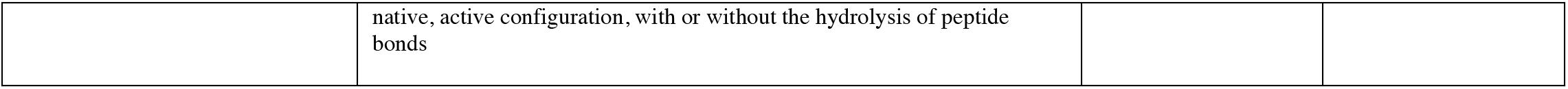
SNP sets significantly associated with autism. The top 25 sets meeting the FDR threshold for significance in either the Broad or CHOP data (N = 102) are shown ranked by q-value (with p-values shown in parenthesis). For each set, q-values shown in italics indicate the set met the significance threshold for that dataset with the most significance q-value shown in bold. The complete set of significant gene sets are provided online in Supplementary Table 1.

To evaluate potential methodological biases, we looked for correlation between the level of significance (p-value) and the number of SNPs per set. The largest correlation found was r^2^ = 0.03, with all other correlations were similarly near zero (Table 2; Supplementary Figure 1). This lack of correlation indicated that differences in the sizes of our sets were not related to differences in the observed levels of significance, and eliminated the possibility of spurious results among the set of significant biological concepts.

Given the limited overlap of genomic coverage between the two studies (Table 1), we were not surprised to find no overlap in the sets identified as significant. The CHOP data yielded the highest number of significant sets with 53 passing our adjusted p-value threshold, while only 15 sets passed the p-value threshold using the Broad data set. Close inspection of these sets showed virtually no overlap in coverage of SNPS between the Broad and CHOP data sets (Figure 1). In other words, when a set was identified as significant using the Broad Affymetrix array the SNPs in the set were not represented on the Illumina array, and vice versa. This suggested that the lack of agreement between the two data sets was due only to experimental design rather than systematic bias associated with our method.

**Figure 1.**
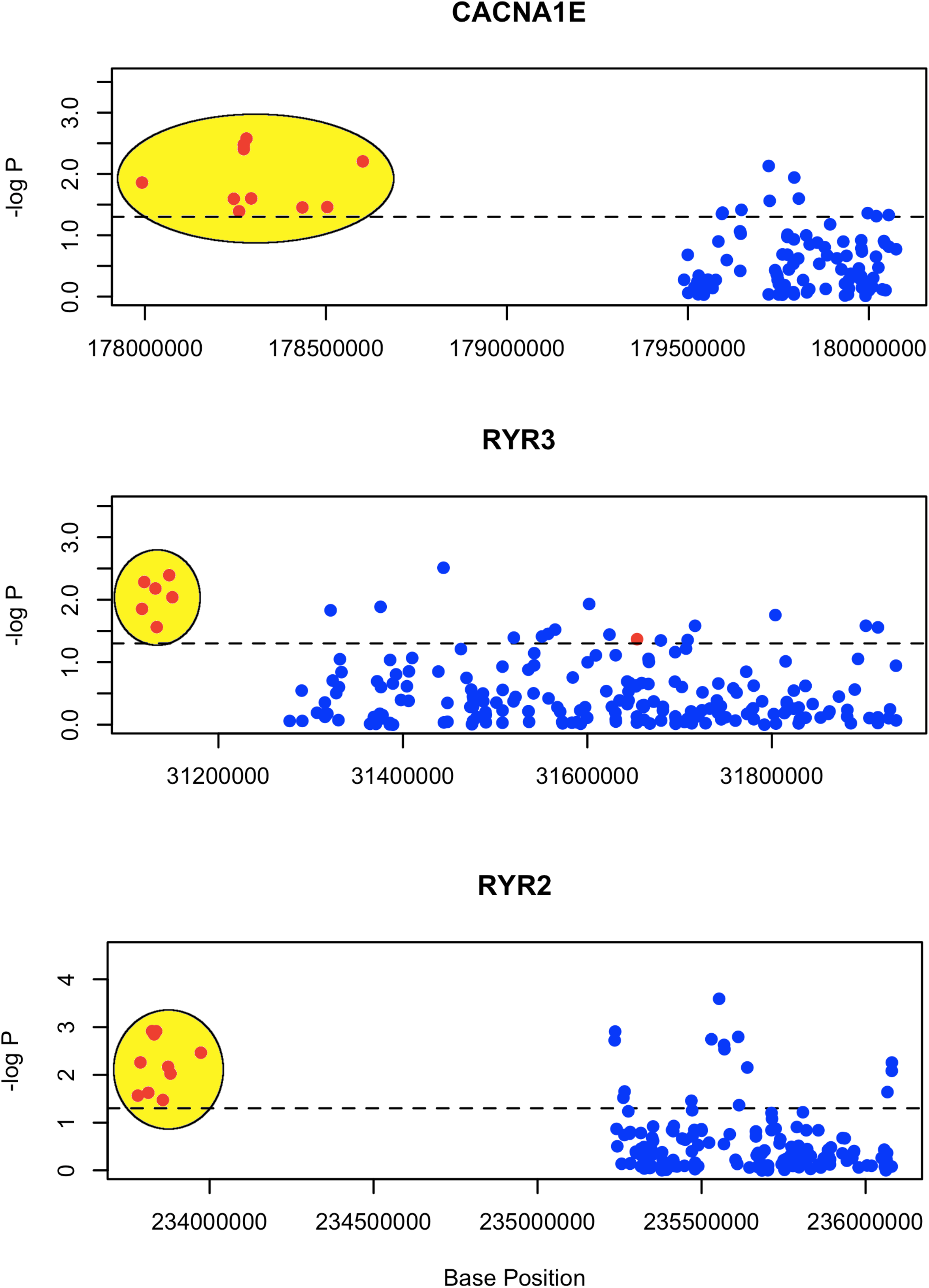
Limited overlap in SNP coverage in the Broad and CHOP data. Provide here is an example of SNP set, “Calcium Channel Activity,” that achieved significance using the CHOP data (red) but not using the Broad data (blue). As depicted, the overlap in SNPs was limited, confirming that the lack of replication was due to experimental design and not biases associated with our enrichment technique.

To characterize redundancy of genes across pathways, we clustered the 68 significant sets according to genes contained in each set. By creating binary profiles of presence and absence of the constituent genes from each set we could compute a pairwise correlation matrix and generate a clustering for all 68 significant sets. Nineteen distinct clusters could be identified including several involved in cell adhesion, immune response, and cancer progression (Figure 2).

**Figure 2.**
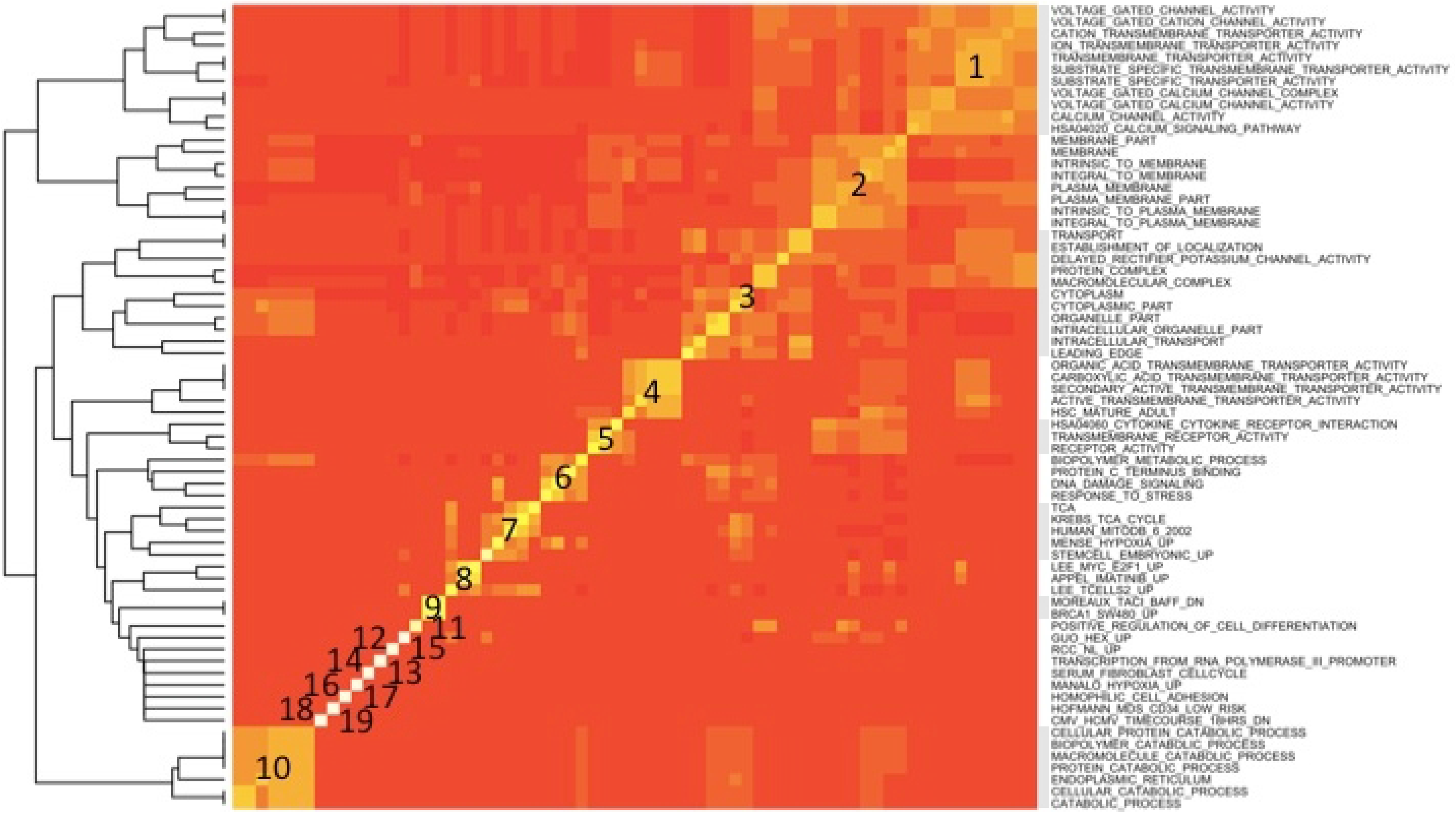
Clustering of significant SNP sets. Gene profiles were constructed for each of the significant sets based on the presence or absence of genes. The profiles were then used to generate the correlation matrix for the SNP sets that in turn was used to perform clustering of the sets, resulting in 19 unique clusters as indicated in the figure. The major themes represented among these 19 clusters were cell adhesion, cancer progression, and immune system response.

### mRNA Expression differences in autistic cases

To validate the significant gene sets and their potential importance to autism, we used published gene expression data from a study of blood-based transcriptional profiles in 17 early-onset autistic cases [17]. Our objective was to determine whether any of the 19 clusters (Figure 2) contained genes with significantly different patterns of expression in autistic individuals when compared with controls. For this, we elected to examine the set in each of the 19 clusters containing the lowest average p value (Table 4). All but two processes contained at least some genes with significant (FDR-adjusted) differential expression, and several had over 90% of its constituent genes exhibiting significant differential expression in autistic cases when compared to controls. In sum, the biological concepts identified as significantly associated with autism genome-wide data appeared to be predictive of differential expression of the same genes in whole-blood samples from a different population of autistic individuals.

**Table 4.**
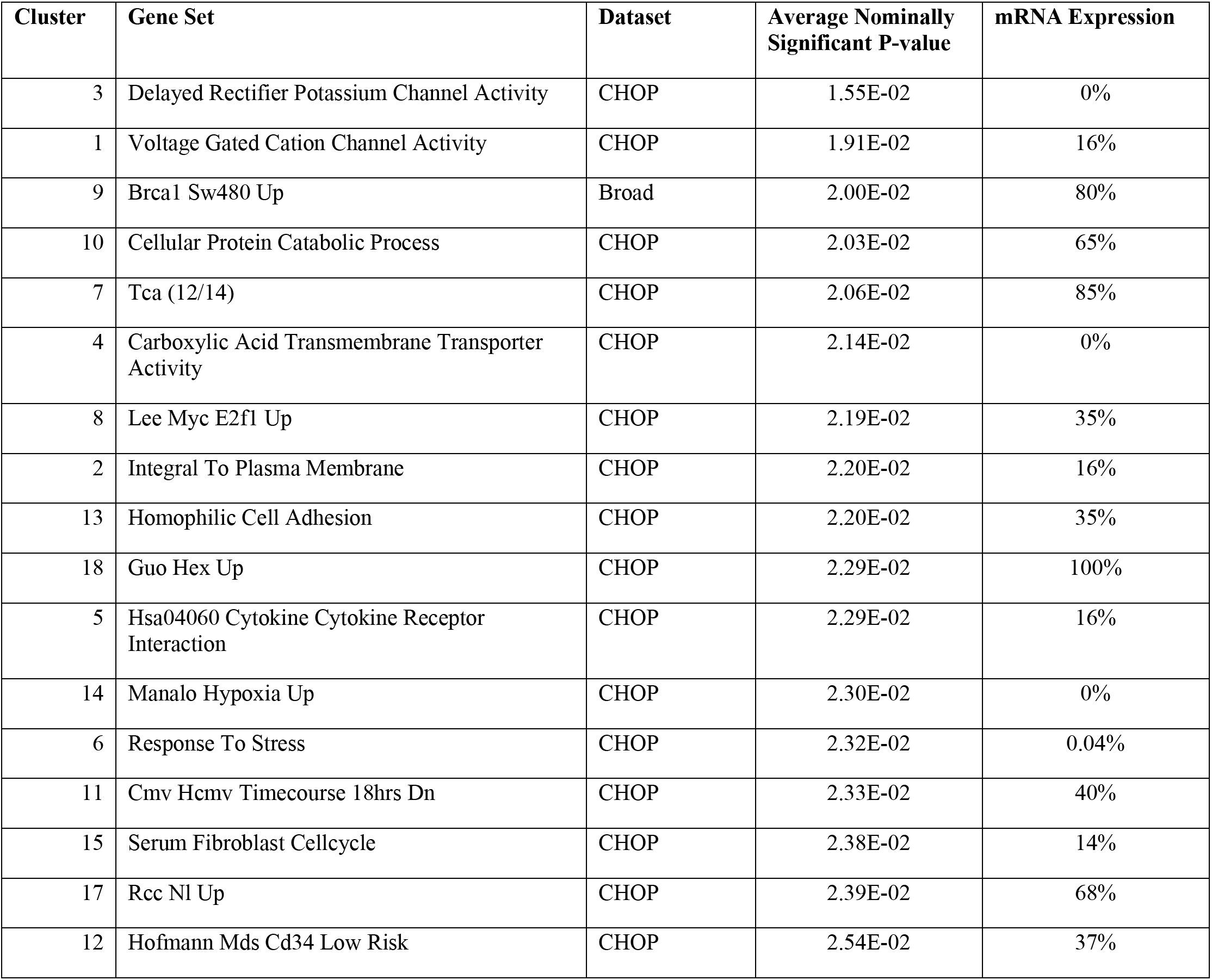
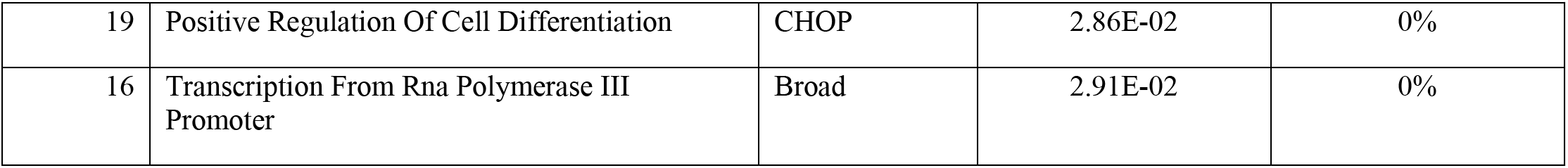
Representative biological concepts among the highest scoring autism gene sets. For each of the 19 clusters revealed through the analysis of the 68 autism-associated sets (Figure 2), we selected the gene set containing the lowest average pvalue as representative, and validated these 19 using both measurements of gene expression (mRNA expression) and tests for enrichment in neurological disease (Table 5). mRNA expression indicates the percentage of genes in the set that have significant, FDR-adjusted p values.

### Neurological disease enrichment

As a second strategy for validation of the significant gene sets, we determined the extent to which each set was enriched for genes involved in autism-related disorders, following from previous work on cross-disease comparison of neurological disorders [18]. Autism spectrum disorders share behaviors and potentially genetic mechanisms with a host of other neurological disorders [18, 19]. Using Autworks, a knowledge base of genetic associations for autism and related conditions, we computed a neurological disease enrichment (NDE) score to determine if our top ranked gene sets contained unusually high percentages of neurological disorder candidates (Table 5). Of the 37 neurological disorders present in Autworks, 36 of them had at least one candidate gene that was a member of one or more of the top-ranking gene sets passing test correction (Table 5). All but three of the 19 representative clusters were significantly enriched (p < 0.05) for known neurological disorder candidate genes.

**Table 5.**
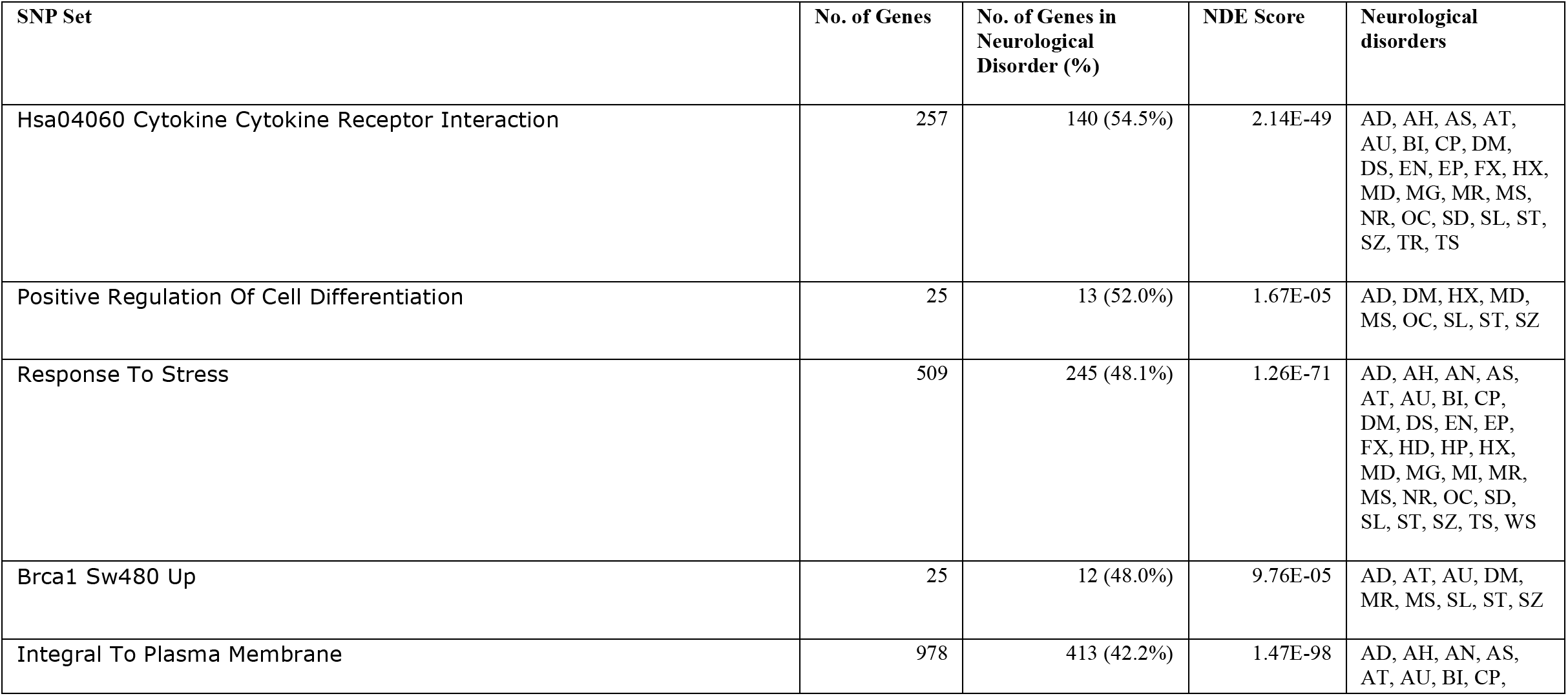
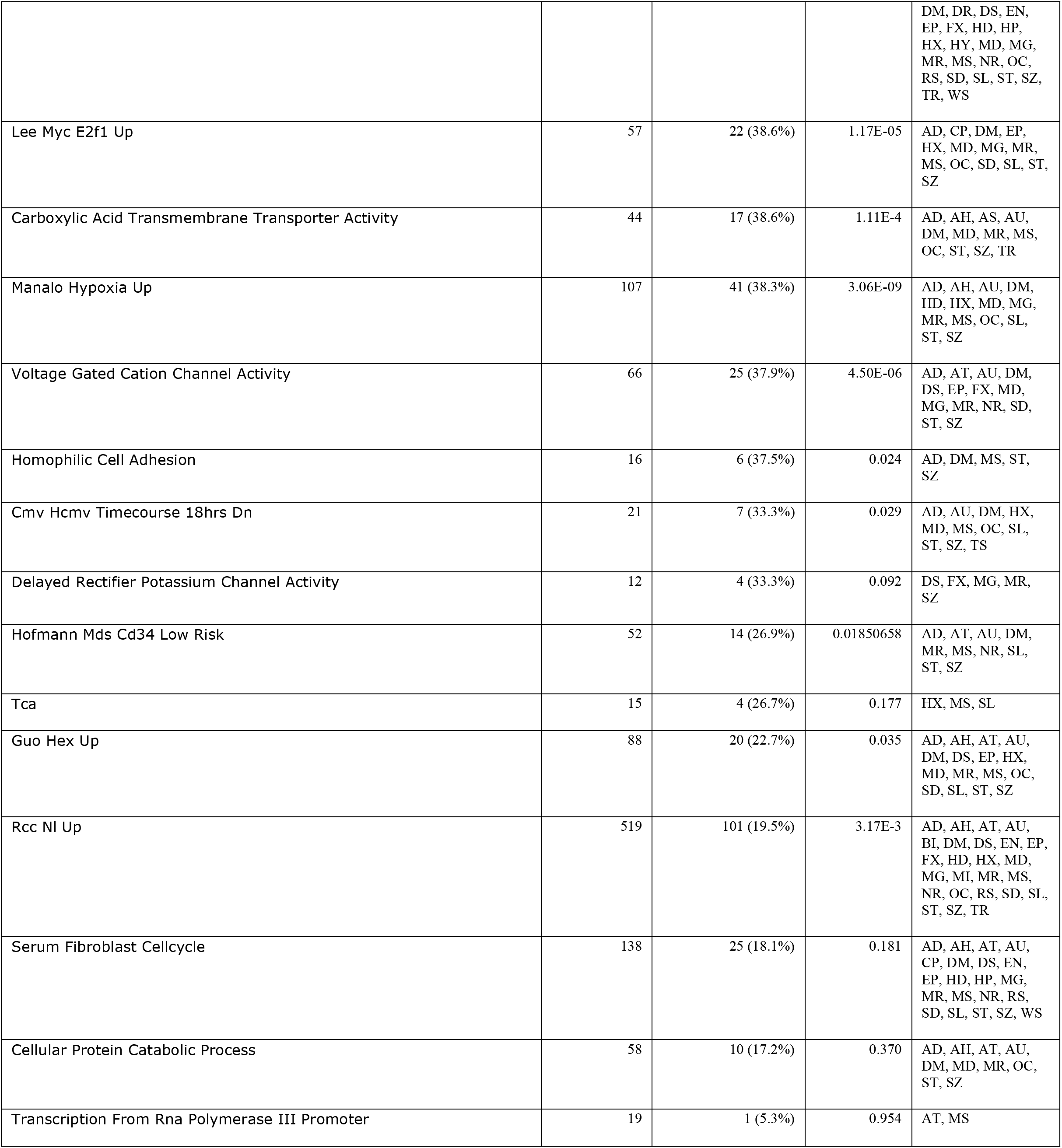
Neurological disease enrichment of autism-associated gene sets. The Autworks resource (autworks.hms.harvard.edu) was used to find genes among the top scoring gene sets that have been implicated in other neurological disorders. A total of 36 conditions were found to share a linked gene with at least one of the significant sets. Significant sets are shown ranked by the proportion of genes implicated in at least one of the 36 conditions (abbreviations provided below). AD=Alzheimer disease; AH=Attention deficit hyperactivity disorder; AN=Angelman Syndrome; AS=Asperger Syndrome; AT=Ataxia; AU=Autism; BI=Brain Injury; CP=Cerebral Palsy; DM=Dementia; DR=De Morsier's Syndrome; DS=Down Syndrome; EN=Encephalopathy; EP=Epilepsy; FX=Fragile X; HD=Huntington Disease; HP=Hypotonia; HX=Hypoxia; HY=Hydrocephalus; MD=Major Depression; MG=Migraine; MI=Microcephaly; MR=Mental Retardation; MS=Multiple Sclerosis; NM=Neuronal Migration Disorders; NR=Neurotoxicity; OC=Obsessive Compulsive Disorder; RS=Rett Syndrome; SD=Seizure Disorder; SL=Systemic Lupus Erythematosus; SP=Spasticity; ST=Stroke; SZ=Schizophrenia; TR=Tourette Syndrome; TS=Tuberous Sclerosis; WL=Williams Syndrome; WS=West Syndrome

## Discussion

The challenge of identifying and characterizing the genetic predisposition to autism has been exacerbated by the low signal-to-noise ratio inherent to genome-wide association studies. In complex disorders, where it is expected that many variants of mild effect induce the disease state, the sample sizes necessary to achieve sufficient statistical power is especially problematic. In fact, the most promising candidate variants in recently published GWAS of autism have detected variants with maximum odds ratios of 0.77 [20] and 1.19 [3], stressing the need for alternative analytical strategies. Here we leveraged prior biological knowledge to supplement standard analysis of GWAS data in order to maximize signal relevant to the genetic basis of autism.

Through application of our knowledge-driven approach (Figure 3), we were able to detect 68 biological concepts significantly associated with autism cases in two studies, one from the Broad Institute (Broad) and the other from the Children’s Hospital of Philadelphia (CHOP). Upon inspection of the genes represented in the 68 concepts of interest, it became apparent that many of the member genes functioned in multiple concepts. Due to this overlap, we were able to cluster the concepts into 19 groups (Figure 2) and identify that the majority of these gene sets could be mapped to three major themes, cell adhesion, cancer response, and immune response.

**Figure 3.**
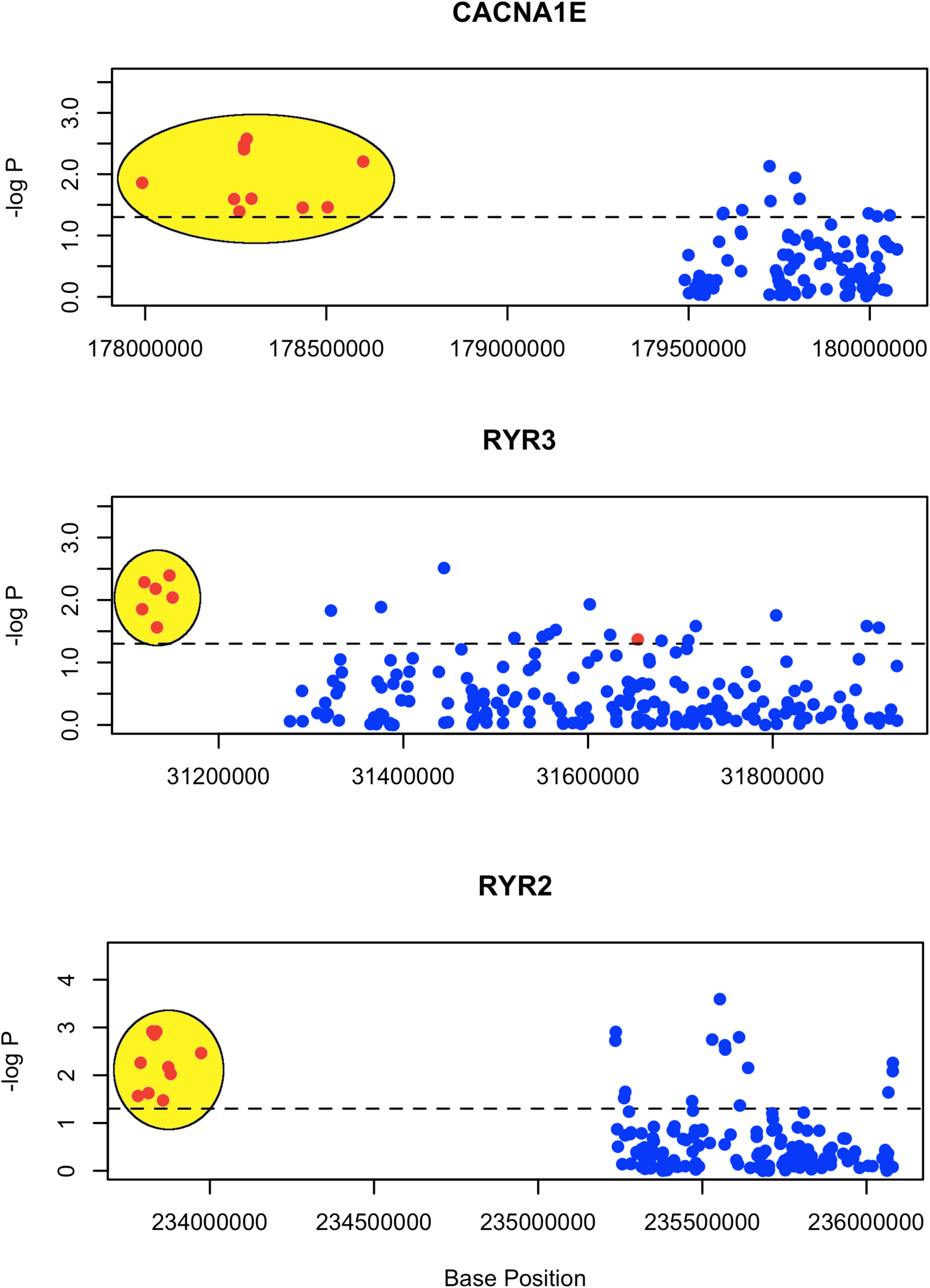
Overview of analytical strategy. First, SNP sets were created by mapping gene sets to the corresponding SNPs using the annotation files provided by the platform vendor. P-values for SNPs were then computed using standard protocols for testing association of genotypes with affection status. After all p-values were calculated, the number of nominally significant SNPs per set was counted. Disease labels were then randomly permuted for N=1000 iterations. Upon each shuffling of the labels, p-values were recomputed and the number of nominally significant SNPs per set was recounted. After the final shuffling of the disease labels, the expected number of nominally significant SNPs was computed for each set given the distribution of the respective counts in the permuted data. Finally, a 2×2 contingency table was constructed for each set and used to compute the Chi Square statistic and p-value for the set.

In strong support of their roles in autism, all but five of our top ranking SNP collections proved to be predictive of significant changes in gene expression in a different study of autistic individuals [17], and all but three yielded significant neurological disease enrichments scores, indicating that they were strongly enriched with candidates linked to disorders already known to be related to autism [18]. These two independent avenues for validation of our sets strongly supported their role in the molecular pathology of autism and stressed the importance of follow-up studies to further refine our understanding of which pathways, and which SNPs in those pathways, may contribute to the manifestation of this disorder.

The 68 significant SNP sets naturally clustered into 19 groups that could be reliably mapped to three themes – cell adhesion, cancer progression, and immune system response. The enrichment of cell adhesion-related concepts in our analysis supports previous results implicating the role of improper cohesive states in the manifestation of autism, among the most recent being the identification of a genome-wide significant variant in the intergenic region between CDH9 and CDH10 [3]. Both CDH9/10 are members of the cadherin superfamily, which includes integral membrane proteins that mediate calcium-dependent cell-cell adhesion. The top scoring of our cell-adhesion concepts was HOMOPHILIC_CELL_ADHESION, a collection of 16 genes that function in the attachment of two identical adhesion molecules in adjacent cells. This set was highly enriched for neurological disorders (Table 4), including three member genes (CADM1, ROBO1, ROBO2) having previous associations with autism [21, 22]. Furthermore, genes in these sets play known roles in central nervous system (CNS) related processes, including activated T cell proliferation [23], axon guidance [24], axon guidance receptor activity [25], axonal fasciculation [26], brain developmental processes [27–29], central nervous system development [25], myelination [26], nervous system development [30], neurite outgrowth [31], neuromuscular junction [32], and positive regulation of axonogenesis [26, 33]. Interestingly, axonal fasciculation is known to be affected by MECP2 levels in Rett Syndrome [34], one of the broad-spectrum autistic disorders. In addition to involvement in the CNS, 6 of the cell adhesion genes identified by our analysis also function in the immune system, including in B cell differentiation [35], defense response [36], immune response [37], leukocyte cell-cell adhesion [35], positive regulation of cytokine secretion [23], positive regulation of immunoglobulin mediated immune response [38], positive regulation of natural killer cell mediated cytotoxicity [39, 40], susceptibility to natural killer cell mediated cytotoxicity [23, 40, 41], susceptibility to T cell mediated cytotoxicity [42], T-cell activation [43, 44], T cell mediated cytotoxicity [23], type 1 fibroblast growth factor receptor binding [45], and synapse development [46]. This intersection in gene function between cell adhesion, CNS development, and immune function provides tantalizing insight into the potential importance of these biological themes in the pathology of autism.

Little evidence has been found to support the role of cancer-related genes in ASDs. However, the presence of several cancer-related concepts among our top scoring and validated SNP sets warranted a deeper look at the function of constituent genes. The top scoring cancer response related concept was BRCA1_SW480_UP (Broad (ALL), mean p = 0.020), a collection of 25 genes affected by BRCA1 expression in breast and lung cancer cell lines that cause dysregulation of the cell cycle and DNA repair processes [47]. Of the 21 genes with SNPs in the Broad data, one (MET) has previously been listed as a likely contributor to autism [48] and 18 of the remaining 20 (90%) have been implicated in at least one other closely related neurological disorder.

MET encodes a proto-oncogene that is involved in cell-cell adhesion, CNS development, and neuron migration. Variants in the MET promoter region that interfere with normal gene transcription have previously been identified in subsets of autism cases [49, 50]. Another gene member gene in this concept, CTNNA1, functions with multiple proteins involved in construction and maintenance of the extracellular matrix. CTNNA1 binds CDH1, a member of the cadherin superfamily that also interacts with MET directly to provide support necessary to promote strong cell-cell adhesive states. The direct interaction of CTNNA1 and CDH1, especially in light of the knowledge that CTNNA1 has previously been linked to both Rett Syndrome and Schizophrenia, may be another mechanism promoting unhealthy cell-cell adhesive states in autism. A third cancer-related gene that promotes healthy cohesive states is the tissue inhibitor of metallopeptidase 1 (TIMP1). The TIMP family of proteins regulates the activity of matrix metalloproteinases (MMPs), a family of peptidases involved in degradation of the extracellular matrix [51]. With extracellular degradation being essential in the routine maintenance and repair of tissue after injury, the appropriate regulation of MMP activity again emphasizes the importance of proper connective states in normal tissue function. Additionally, the balance of TIMP1/MMP9 levels is thought to be important in blood-brain barrier (BBB) function [52]. TIMP1 levels have previously been shown to be dysregulated in the serum and cerebrospinal fluid (CSF) of patients with neurological disorders related to autism [53–57], suggesting improper TIMP1/MMP9 levels and dysfunction of the BBB may contribute to the onset and progression of neurological disorders.

The role of the immune system in the etiology of ASDs has been widely suspect, particularly the plausibility of adverse events in the autoimmune system [58, 59]. Accordingly, the presence of several immune-related concepts among the 19 concept classes was particularly intriguing. The immune-related concept showing the highest associated was HSA04060_CYTOKINE_CYTOKINE_RECEPTOR_INTERACTION, a set of 257 genes involved in cytokine-cytokine receptor interaction [60]. Of the 257 genes, 15 (CCL2, CD40LG, EGF, GHR [61], HGF, IFNG [62], IL18R, LEP, LEPR, MET [48–50, 63], PRL [64], PRLR [64], TGFB1, TNF [65], TPO) already have been labeled as candidate genes for autism, and 202 have some association with neurological disorders that are linked to autism.

We have demonstrated that knowledge-driven approaches to signal detection can detect autism-related signal previously missed by alternative approaches. And, we have demonstrated that two of the largest published GWAS data sets on autism to-date contain an abundance of significant pathways with SNPs that are highly likely to play linked to the molecular pathology. These pathways could be largely verified in two ways, first by transcriptional profiles of autistic individuals, and second by their enrichment of neurological disease candidates. In the former, we were able to demonstrate that all but 5 of the 19 most significant pathways contained genes that are significantly differentially expressed in autistic individuals. In the latter, leveraging published studies on autism-related disorders, we were able to demonstrate that all but three of these 19 pathways contained significant percentages of genes linked to neurological disease.

Our reanalysis of autism genome-wide association data identified significant allelic differences among autistic individuals that correspond to a limited number of biological concepts. While it remains difficult to determine which SNPs among these concepts are the most informative to the genetic bases of autism, our results point to a highly promising set of candidates that are worthy of further investigation.

## Conclusions

Our approach identified significant signal in two historical autism GWAS datasets. The genomic variants were enriched in autism-related neurological disorders and linked to genes that were significantly differentially expressed in autistic individuals when compared to controls. These genetic variants represent a high priority set of candidates worthy of deeper inspection in a larger cohort of autistic individuals. In sum, our work provides exciting new leads into the genetic underpinnings of autism and highlights the importance of reanalysis of genomic studies of complex disease using prior knowledge of genetic organization.

## Materials and Methods

### Ethics Statement

Our study (number: M18096-101) has been evaluated by the Institutional Review Board and identified exempt as defined under 45CFR46.102(f) and as meeting the conditions regarding coded biological specimens or data. As such, (a) the specimens/data were not collected specifically for the research through an interaction or intervention with a living person, and (b) the investigators cannot “readily ascertain” the identity of the individual who provided the specimen/data to whom any code pertains.

**{Bailey, 1995 #380}**

### Genotype samples

Genotyping data were acquired from the Autism Genetic Resource Exchange (AGRE) [66], consisting of two previously described cohorts from the Broad Institute (Broad) [2] and the Children’s Hospital of Philadelphia (CHOP) [44]. The Broad cohort consisted of 2,883 subjects from 751 families genotyped at 500K SNPs with probands diagnosed using the Autism Diagnostic Interview-Revised (ADI-R) [67], and the CHOP cohort consisted of 4,444 subjects from 943 families genotyped at 550K SNPs with probands diagnosed using both the ADI-R and the Autism Diagnostic Observation Schedule (ADOS) [68].

Standard quality control measures were applied using PLINK [69]. SNPs were excluded from the analysis if minor allele frequency < 0.05, call rate < 0.80, or Hardy-Weinberg Equilibrium (HWE) was not met (p < 0.01). Additionally, genotypes for all nuclear family members were set to “missing” for SNPs in violation of Mendelian rules of inheritance. Genotyping samples were removed where sex or affected status were unknown, or genotyping call rate < 0.80. Finally, subjects of European descent were selected for further analysis, leaving 727 families containing 3,322 subjects and 557 families containing 2,137 subjects for the CHOP and Broad cohorts, respectively (Table 1). After applying quality control measures and removing SNPs not required for the set-based analysis, 269,214 (ALL) and 112,403 (PRI) SNPs remained for the Broad cohort, and 264,543 (ALL) and 102,194 (PRI) SNPs remained for the CHOP cohort (Table 1).

### SNP set construction

SNP identifiers were mapped to the 1,892 C2 and 1,454 C5 MSigDB [70] gene sets based on the SNP-to-gene associations specified in the Affymetrix 5.0 and Illumina HumanHap550 genotyping platform annotation files for cohorts from the Broad Institute [2] (Broad) and Children’s Hospital of Philadelphia [3] (CHOP), respectively. All SNPs listed as associated with a gene were mapped to the set.

### Set-based analysis

We devised an analytical strategy that incorporates *a priori* biological knowledge to collectively test for significant association of groups of SNPs with autism in the genome-wide association data utilized here. The outline of our strategy is depicted in Figure 3 and described below.

#### 1. Construction of SNP sets

SNP identifiers were mapped to the C2 (curated gene collections) and C5 (gene sets from the biological process ontology of Gene Ontology) MSigDB [70] gene sets based on the SNP-gene relationships specified in the respective genotyping platform annotation files to create SNP sets representing the biological concept of the original gene set.

#### 2. Compute SNP observed p-values and set counts

For each SNP S_i_, a p-value, 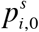, was calculated following the procedure previously described [16] as appropriate for family-based cohorts; an additive effects model was used to code the genotypes. The number of SNPs in each SNP set SS_j_ with 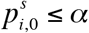, 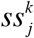, was then counted and saved for later use.

#### 3. Compute SNP permuted p-values and set counts

Affection status was randomly assigned in children for K = 1000 iterations, keeping the ratio of affected to unaffected individuals constant. After each reassignment of affection status, p-values were calculated for each S_i_, with 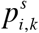 representing the p-value computed for the i^th^ SNP at the k^th^ permutation. The number of SNPs in each SNP set SS_j_ with 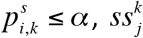, was then counted for each k ∈ [1, K]. The 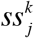 for k > 0 generate an appropriate null distribution for each SS_j_ since they preserve the original biological concept as well as account for the correlation structure inherently present in the genotyping data.

#### 4. Assessing SNP set significance

Once all 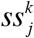 have been computed, the expected value of *ss*_*j*_ can be computed as:

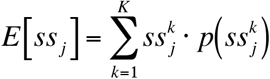

Sets with *E*[*ss*_*j*_] ≤ 5 or 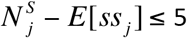, with 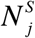 representing the number of SNPs in set j, were excluded from further consideration due to the low expected value compromising the reliability of the test statistic calculation.

To assess the significance of each 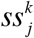, p-values for each SNP set were calculated as follows:

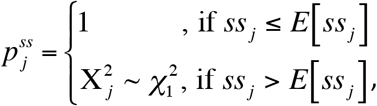

where 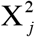 is the Pearson’s *χ*^2^ test statistic for the j^th^ set computed from the contingency table:

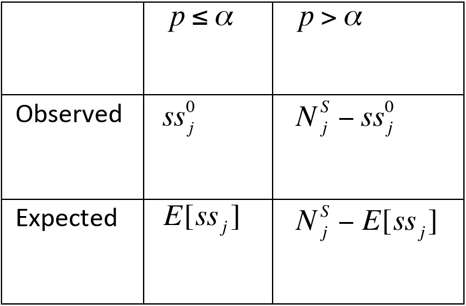

Finally, q-values [71] for each set were computed using the qvalue package in R and all SNP sets with *q*_*j*_ ≤ *q* were considered significant.

### Clustering of significant sets

Once significant sets of SNPs were identified, we clustered the sets to reduce redundancy of the representative genes and improve our ability to identify which major biological themes were enriched among our autism cases. To do so, we created binary profiles of genes present or absent in each of the sets passing our significance cutoff. Then, we computed a Pearson correlation coefficient for all pairs of binary profiles to create a pairwise correlation matrix that could be used to generate a simple tree to visualize clustering among the sets.

### Neurological disease enrichment (NDE) scores

To assess the representation of neurologically related genes in each of the biological concepts, we computed a neurological disease enrichment (NDE) scores as follows:

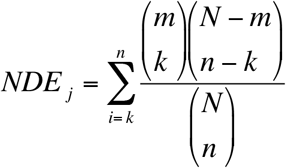

where N = 23402, the number of human gene entries in Entrez gene (http://www.ncbi.nlm.nih.gov/gene) at the time of this writing, m = 3511, the total number of genes with known association to a neurological disorder, n is the number of genes in gene set j, and k is the number of neurologically related genes observed in gene set j. Intuitively, this represents the probability of a biological concept containing at least as many neurological disorder genes as was observed. Using Autworks, a knowledge base of genetic associations for autism and related conditions, we computed a neurological disease enrichment (NDE) score, to determine if our top ranked gene sets contained unusually high percentages of neurological disorder candidates (Table 5).

### mRNA expression data processing

We downloaded GSE6575 [17, 59] from the Gene Expression Omnibus (GEO) for validation of the sets identified by our SNP-based gene enrichment method. This dataset consisted of 17 samples of autistic patients without regression, 18 patients with regression, 9 patients with mental retardation or developmental delay, and 12 typically developing children from the general population. In this cohort, total RNA was extracted from whole blood samples using the PaxGene Blood RNA System and run on Affymetrix U133plus2.0. For the purposes of our study, we compared the 17 autistic cases without regression with the 12 control samples from the general population using a student’s t-test to generate nominal p values for every transcript. The 18 autistic cases with regression had previously been shown to contain limited signal [18] and were excluded from the validation steps in this study. Preprocessing and expression analyses were done with the Bioinformatics Toolbox Version 2.6 (For Matlab R2007a+). GCRMA was used for background adjustment and control probe intensities were used to estimate non-specific binding [72]. Housekeeping genes, gene expression data with empty gene symbols, genes with very low absolute expression values, and genes with a small variance across samples were removed from the preprocessed dataset. To adjust for multiple testing, we used the q-value calculation [71], a measurement framed in terms of the false discovery rate [73], considering q < 0.05 an indication of significant differential expression.

## List of abbreviations

GWAS: genome-wide association study
SNP: single-nucleotide polymorphism
CHOP: Children’s Hospital of Philadelphia

## Competing interests

None

## Authors’ contributions

DPW conceived the project, designed the algorithms, participated in the analysis, and wrote the manuscript. JYJ, FJE, PJT participated in interpretation of the results and edited the manuscript.

## Acknowledgements

We would like to thank Badri Vadarajan, Vincent Fusaro and members of the Tonellato-Wall weekly lab meetings for vital input on the computational approach and interpretation of results. We also thank the Autism Genetic Research Exchange for providing access to data, the families enrolled in the AGRE project for participating in this important endeavor, and Vlad Kustanovich for assistance with downloading and handling AGRE data. Finally, we acknowledge the Translational Research Program at Children’s Hospital of Boston for support.

---

**Supplementary Figure 1.**
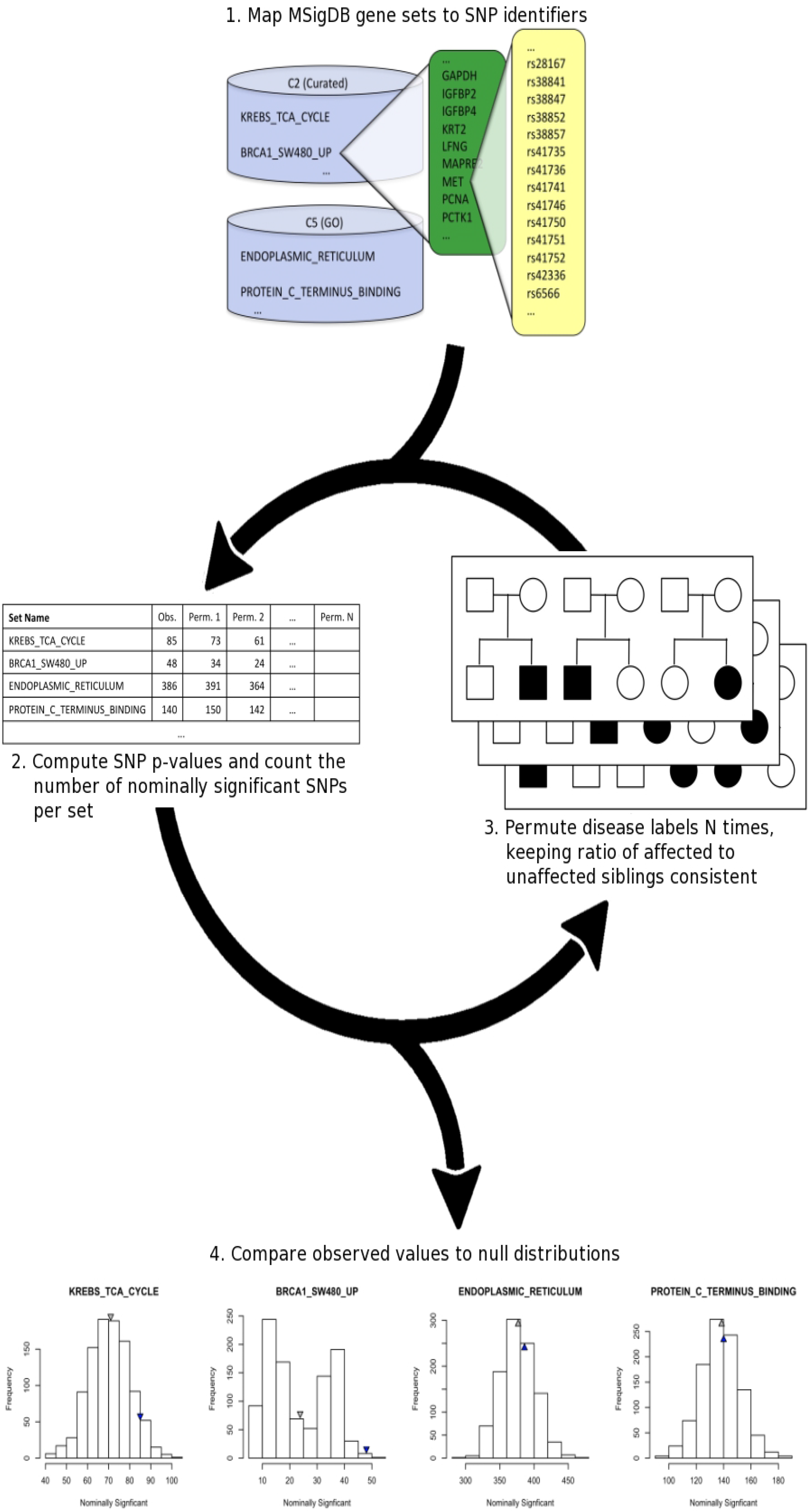
Gene set size versus p-value. Correlation coefficients were calculated and correlation plots generated to evaluate any potential bias arising from SNP set size. No substantial correlation was found between set size and p-value (maximum r^2^ = 0.0349), even with larger sets having higher statistical power to detect subtle differences. Points for sets passing FDR correction are shown in red. We examined correlation for all SNPs annotated to a gene and only SNPs within the primary coding transcript (PRI) for both Broad and CHOP data sets.

## Notes

### Competing Interest Statement

The authors have declared no competing interest.

### Summary of Updates

To add additional references.

## References

1. Freitag CM: The genetics of autistic disorders and its clinical relevance: a review of the literature. Mol Psychiatry 2007, 12(1):2–22.

2. Weiss LA, Shen Y, Korn JM, Arking DE, Miller DT, Fossdal R, Saemundsen E, Stefansson H, Ferreira MA, Green T et al.: Association between microdeletion and microduplication at 16p11.2 and autism. N Engl J Med 2008, 358(7):667–675.

3. Wang K, Zhang H, Ma D, Bucan M, Glessner JT, Abrahams BS, Salyakina D, Imielinski M, Bradfield JP, Sleiman PM et al.: Common genetic variants on 5p14.1 associate with autism spectrum disorders. Nature 2009, 459(7246):528–533.

4. Goldstein DB: Common genetic variation and human traits. N Engl J Med 2009, 360(17):1696–1698.

5. Kraft P, Hunter DJ: Genetic risk prediction--are we there yet? N Engl J Med 2009, 360(17):1701–1703.

6. Kraft P, Wacholder S, Cornelis MC, Hu FB, Hayes RB, Thomas G, Hoover R, Hunter DJ, Chanock S: Beyond odds ratios--communicating disease risk based on genetic profiles. Nat Rev Genet 2009, 10(4):264–269.

7. Hirschhorn JN: Genomewide association studies--illuminating biologic pathways. N Engl J Med 2009, 360(17):1699–1701.

8. Baranzini SE, Galwey NW, Wang J, Khankhanian P, Lindberg R, Pelletier D, Wu W, Uitdehaag BM, Kappos L, Polman CH et al.: Pathway and network-based analysis of genome-wide association studies in multiple sclerosis. Hum Mol Genet 2009, 18(11):2078–2090.

9. Elbers CC, van Eijk KR, Franke L, Mulder F, van der Schouw YT, Wijmenga C, Onland-Moret NC: Using genome-wide pathway analysis to unravel the etiology of complex diseases. Genet Epidemiol 2009, 33(5):419–431.

10. Holmans P, Green EK, Pahwa JS, Ferreira MA, Purcell SM, Sklar P, Owen MJ, O’Donovan MC, Craddock N: Gene ontology analysis of GWA study data sets provides insights into the biology of bipolar disorder. Am J Hum Genet 2009, 85(1):13–24.

11. Medina I, Montaner D, Bonifaci N, Pujana MA, Carbonell J, Tarraga J, Al-Shahrour F, Dopazo J: Gene set-based analysis of polymorphisms: finding pathways or biological processes associated to traits in genome-wide association studies. Nucleic Acids Res 2009, 37(Web Server issue):W340–344.

12. O’Dushlaine C, Kenny E, Heron EA, Segurado R, Gill M, Morris DW, Corvin A: The SNP ratio test: pathway analysis of genome-wide association datasets. Bioinformatics 2009, 25(20):2762–2763.

13. Peng G, Luo L, Siu H, Zhu Y, Hu P, Hong S, Zhao J, Zhou X, Reveille JD, Jin L et al.: Gene and pathway-based second-wave analysis of genome-wide association studies. Eur J Hum Genet 2010, 18(1):111–117.

14. Torkamani A, Topol EJ, Schork NJ: Pathway analysis of seven common diseases assessed by genome-wide association. Genomics 2008, 92(5):265–272.

15. Wang K, Li M, Bucan M: Pathway-Based Approaches for Analysis of Genomewide Association Studies. Am J Hum Genet 2007, 81(6).

16. Laird NM, Lange C: Family-based designs in the age of large-scale gene-association studies. Nat Rev Genet 2006, 7(5):385–394.

17. Gregg JP, Lit L, Baron CA, Hertz-Picciotto I, Walker W, Davis RA, Croen LA, Ozonoff S, Hansen R, Pessah IN et al.: Gene expression changes in children with autism. Genomics 2008, 91(1):22–29.

18. Wall DP, Esteban FJ, Deluca TF, Huyck M, Monaghan T, Velez de Mendizabal N, Goni J, Kohane IS: Comparative analysis of neurological disorders focuses genome-wide search for autism genes. Genomics 2009, 93(2):120–129.

19. Belmonte MK, Bourgeron T: Fragile X syndrome and autism at the intersection of genetic and neural networks. Nat Neurosci 2006, 9(10):1221–1225.

20. Weiss LA, Arking DE, Daly MJ, Chakravarti A: A genome-wide linkage and association scan reveals novel loci for autism. Nature 2009, 461(7265):802–808.

21. Anitha A, Nakamura K, Yamada K, Suda S, Thanseem I, Tsujii M, Iwayama Y, Hattori E, Toyota T, Miyachi T et al.: Genetic analyses of roundabout (ROBO) axon guidance receptors in autism. Am J Med Genet B Neuropsychiatr Genet 2008, 147B(7):1019–1027.

22. Zhiling Y, Fujita E, Tanabe Y, Yamagata T, Momoi T, Momoi MY: Mutations in the gene encoding CADM1 are associated with autism spectrum disorder. Biochem Biophys Res Commun 2008, 377(3):926–929.

23. Boles KS, Barchet W, Diacovo T, Cella M, Colonna M: The tumor suppressor TSLC1/NECL-2 triggers NK-cell and CD8+ T-cell responses through the cell-surface receptor CRTAM. Blood 2005, 106(3):779–786.

24. Shu T, Richards LJ: Cortical axon guidance by the glial wedge during the development of the corpus callosum. J Neurosci 2001, 21(8):2749–2758.

25. Kidd T, Brose K, Mitchell KJ, Fetter RD, Tessier-Lavigne M, Goodman CS, Tear G: Roundabout controls axon crossing of the CNS midline and defines a novel subfamily of evolutionarily conserved guidance receptors. Cell 1998, 92(2):205–215.

26. Kuja-Panula J, Kiiltomaki M, Yamashiro T, Rouhiainen A, Rauvala H: AMIGO, a transmembrane protein implicated in axon tract development, defines a novel protein family with leucine-rich repeats. J Cell Biol 2003, 160(6):963–973.

27. Stagi M, Fogel AI, Biederer T: SynCAM 1 participates in axo-dendritic contact assembly and shapes neuronal growth cones. Proc Natl Acad Sci U S A 2010, 107(16):7568–7573.

28. Galuska SP, Rollenhagen M, Kaup M, Eggers K, Oltmann-Norden I, Schiff M, Hartmann M, Weinhold B, Hildebrandt H, Geyer R et al.: Synaptic cell adhesion molecule SynCAM 1 is a target for polysialylation in postnatal mouse brain. Proc Natl Acad Sci U S A 2010, 107(22):10250–10255.

29. Nguyen Ba-Charvet KT, Brose K, Marillat V, Kidd T, Goodman CS, Tessier-Lavigne M, Sotelo C, Chedotal A: Slit2-Mediated chemorepulsion and collapse of developing forebrain axons. Neuron 1999, 22(3):463–473.

30. Sundaresan V, Roberts I, Bateman A, Bankier A, Sheppard M, Hobbs C, Xiong J, Minna J, Latif F, Lerman M et al.: The DUTT1 gene, a novel NCAM family member is expressed in developing murine neural tissues and has an unusually broad pattern of expression. Mol Cell Neurosci 1998, 11(1-2):29–35.

31. Hagiyama M, Ichiyanagi N, Kimura KB, Murakami Y, Ito A: Expression of a soluble isoform of cell adhesion molecule 1 in the brain and its involvement in directional neurite outgrowth. Am J Pathol 2009, 174(6):2278–2289.

32. van der Flier A, Gaspar AC, Thorsteinsdottir S, Baudoin C, Groeneveld E, Mummery CL, Sonnenberg A: Spatial and temporal expression of the beta1D integrin during mouse development. Dev Dyn 1997, 210(4):472–486.

33. Hivert B, Liu Z, Chuang CY, Doherty P, Sundaresan V: Robo1 and Robo2 are homophilic binding molecules that promote axonal growth. Mol Cell Neurosci 2002, 21(4):534–545.

34. Degano AL, Pasterkamp RJ, Ronnett GV: MeCP2 deficiency disrupts axonal guidance, fasciculation, and targeting by altering Semaphorin 3F function. Mol Cell Neurosci 2009, 42(3):243–254.

35. Ryan DH, Nuccie BL, Abboud CN, Winslow JM: Vascular cell adhesion molecule-1 and the integrin VLA-4 mediate adhesion of human B cell precursors to cultured bone marrow adherent cells. J Clin Invest 1991, 88(3):995–1004.

36. de la Fuente MA, Pizcueta P, Nadal M, Bosch J, Engel P: CD84 leukocyte antigen is a new member of the Ig superfamily. Blood 1997, 90(6):2398–2405.

37. Suzuki K, Hu D, Bustos T, Zlotogora J, Richieri-Costa A, Helms JA, Spritz RA: Mutations of PVRL1, encoding a cell-cell adhesion molecule/herpesvirus receptor, in cleft lip/palate-ectodermal dysplasia. Nat Genet 2000, 25(4):427–430.

38. Bachelet I, Munitz A, Mankutad D, Levi-Schaffer F: Mast cell costimulation by CD226/CD112 (DNAM-1/Nectin-2): a novel interface in the allergic process. J Biol Chem 2006, 281(37):27190–27196.

39. Pende D, Castriconi R, Romagnani P, Spaggiari GM, Marcenaro S, Dondero A, Lazzeri E, Lasagni L, Martini S, Rivera P et al.: Expression of the DNAM-1 ligands, Nectin-2 (CD112) and poliovirus receptor (CD155), on dendritic cells: relevance for natural killer-dendritic cell interaction. Blood 2006, 107(5):2030–2036.

40. Shingai T, Ikeda W, Kakunaga S, Morimoto K, Takekuni K, Itoh S, Satoh K, Takeuchi M, Imai T, Monden M et al.: Implications of nectin-like molecule-2/IGSF4/RA175/SgIGSF/TSLC1/SynCAM1 in cell-cell adhesion and transmembrane protein localization in epithelial cells. J Biol Chem 2003, 278(37):35421–35427.

41. Bottino C, Castriconi R, Pende D, Rivera P, Nanni M, Carnemolla B, Cantoni C, Grassi J, Marcenaro S, Reymond N et al.: Identification of PVR (CD155) and Nectin-2 (CD112) as cell surface ligands for the human DNAM-1 (CD226) activating molecule. J Exp Med 2003, 198(4):557–567.

42. Tahara-Hanaoka S, Shibuya K, Onoda Y, Zhang H, Yamazaki S, Miyamoto A, Honda S, Lanier LL, Shibuya A: Functional characterization of DNAM-1 (CD226) interaction with its ligands PVR (CD155) and nectin-2 (PRR-2/CD112). Int Immunol 2004, 16(4):533–538.

43. Tangye SG, Nichols KE, Hare NJ, van de Weerdt BC: Functional requirements for interactions between CD84 and Src homology 2 domain-containing proteins and their contribution to human T cell activation. J Immunol 2003, 171(5):2485–2495.

44. Wang M, Windgassen D, Papoutsakis ET: Comparative analysis of transcriptional profiling of CD3+, CD4+ and CD8+ T cells identifies novel immune response players in T-cell activation. BMC Genomics 2008, 9:225.

45. Owczarek S, Kiryushko D, Larsen MH, Kastrup JS, Gajhede M, Sandi C, Berezin V, Bock E, Soroka V: Neuroplastin-55 binds to and signals through the fibroblast growth factor receptor. FASEB J 2010, 24(4):1139–1150.

46. Huang Z, Shimazu K, Woo NH, Zang K, Muller U, Lu B, Reichardt LF: Distinct roles of the beta 1-class integrins at the developing and the mature hippocampal excitatory synapse. J Neurosci 2006, 26(43):11208–11219.

47. MacLachlan TK, Somasundaram K, Sgagias M, Shifman Y, Muschel RJ, Cowan KH, El-Deiry WS: BRCA1 effects on the cell cycle and the DNA damage response are linked to altered gene expression. J Biol Chem 2000, 275(4):2777–2785.

48. Abrahams BS, Geschwind DH: Advances in autism genetics: on the threshold of a new neurobiology. Nat Rev Genet 2008, 9(5):341–355.

49. Campbell DB, Sutcliffe JS, Ebert PJ, Militerni R, Bravaccio C, Trillo S, Elia M, Schneider C, Melmed R, Sacco R et al.: A genetic variant that disrupts MET transcription is associated with autism. Proc Natl Acad Sci U S A 2006, 103(45):16834–16839.

50. Sousa I, Clark TG, Toma C, Kobayashi K, Choma M, Holt R, Sykes NH, Lamb JA, Bailey AJ, Battaglia A et al.: MET and autism susceptibility: family and case-control studies. Eur J Hum Genet 2009, 17(6):749–758.

51. Chandler S, Miller KM, Clements JM, Lury J, Corkill D, Anthony DC, Adams SE, Gearing AJ: Matrix metalloproteinases, tumor necrosis factor and multiple sclerosis: an overview. J Neuroimmunol 1997, 72(2):155–161.

52. Rosenberg GA, Estrada EY, Dencoff JE: Matrix metalloproteinases and TIMPs are associated with blood-brain barrier opening after reperfusion in rat brain. Stroke 1998, 29(10):2189–2195.

53. Cuadrado E, Rosell A, Penalba A, Slevin M, Alvarez-Sabin J, Ortega-Aznar A, Montaner J: Vascular MMP-9/TIMP-2 and neuronal MMP-10 up-regulation in human brain after stroke: a combined laser microdissection and protein array study. J Proteome Res 2009, 8(6):3191–3197.

54. Lorenzl S, Albers DS, LeWitt PA, Chirichigno JW, Hilgenberg SL, Cudkowicz ME, Beal MF: Tissue inhibitors of matrix metalloproteinases are elevated in cerebrospinal fluid of neurodegenerative diseases. J Neurol Sci 2003, 207(1-2):71–76.

55. Robak E, Wierzbowska A, Chmiela M, Kulczycka L, Sysa-Jedrejowska A, Robak T: Circulating total and active metalloproteinase-9 and tissue inhibitor of metalloproteinases-1 in patients with systemic lupus erythomatosus. Mediators Inflamm 2006, 2006(1):17898.

56. Suenaga N, Ichiyama T, Kubota M, Isumi H, Tohyama J, Furukawa S: Roles of matrix metalloproteinase-9 and tissue inhibitors of metalloproteinases 1 in acute encephalopathy following prolonged febrile seizures. J Neurol Sci 2008, 266(1-2):126–130.

57. Suryadevara R, Holter S, Borgmann K, Persidsky R, Labenz-Zink C, Persidsky Y, Gendelman HE, Wu L, Ghorpade A: Regulation of tissue inhibitor of metalloproteinase-1 by astrocytes: links to HIV-1 dementia. Glia 2003, 44(1):47–56.

58. Atladottir HO, Pedersen MG, Thorsen P, Mortensen PB, Deleuran B, Eaton WW, Parner ET: Association of family history of autoimmune diseases and autism spectrum disorders. Pediatrics 2009, 124(2):687–694.

59. Gonzalez-Gronow M, Cuchacovich M, Francos R, Cuchacovich S, Del Pilar Fernandez M, Blanco A, Bowers EV, Kaczowka S, Pizzo SV: Antibodies against the voltage-dependent anion channel (VDAC) and its protective ligand hexokinase-I in children with autism. J Neuroimmunol 2010, 227(1-2):153–161.

60. Kanehisa M, Araki M, Goto S, Hattori M, Hirakawa M, Itoh M, Katayama T, Kawashima S, Okuda S, Tokimatsu T et al.: KEGG for linking genomes to life and the environment. Nucleic Acids Res 2008, 36(Database issue):D480–484.

61. Mills JL, Hediger ML, Molloy CA, Chrousos GP, Manning-Courtney P, Yu KF, Brasington M, England LJ: Elevated levels of growth-related hormones in autism and autism spectrum disorder. Clin Endocrinol (Oxf) 2007, 67(2):230–237.

62. Ashwood P, Anthony A, Torrente F, Wakefield AJ: Spontaneous mucosal lymphocyte cytokine profiles in children with autism and gastrointestinal symptoms: mucosal immune activation and reduced counter regulatory interleukin-10. J Clin Immunol 2004, 24(6):664–673.

63. Campbell DB, Warren D, Sutcliffe JS, Lee EB, Levitt P: Association of MET with social and communication phenotypes in individuals with autism spectrum disorder. Am J Med Genet B Neuropsychiatr Genet 2010, 153B(2):438–446.

64. Yrigollen CM, Han SS, Kochetkova A, Babitz T, Chang JT, Volkmar FR, Leckman JF, Grigorenko EL: Genes controlling affiliative behavior as candidate genes for autism. Biol Psychiatry 2008, 63(10):911–916.

65. Guerini FR, Bolognesi E, Manca S, Sotgiu S, Zanzottera M, Agliardi C, Usai S, Clerici M: Family-based transmission analysis of HLA genetic markers in Sardinian children with autistic spectrum disorders. Hum Immunol 2009, 70(3):184–190.

66. Geschwind DH, Sowinski J, Lord C, Iversen P, Shestack J, Jones P, Ducat L, Spence SJ: The autism genetic resource exchange: a resource for the study of autism and related neuropsychiatric conditions. Am J Hum Genet 2001, 69(2):463–466.

67. Lord C, Rutter M, Le Couteur A: Autism Diagnostic Interview-Revised: a revised version of a diagnostic interview for caregivers of individuals with possible pervasive developmental disorders. J Autism Dev Disord 1994, 24(5):659–685.

68. Lord C, Risi S, Lambrecht L, Cook EH, Jr., Leventhal BL, DiLavore PC, Pickles A, Rutter M: The autism diagnostic observation schedule-generic: a standard measure of social and communication deficits associated with the spectrum of autism. J Autism Dev Disord 2000, 30(3):205–223.

69. Purcell S, Neale B, Todd-Brown K, Thomas L, Ferreira MA, Bender D, Maller J, Sklar P, de Bakker PI, Daly MJ et al.: PLINK: a tool set for whole-genome association and population-based linkage analyses. Am J Hum Genet 2007, 81(3):559–575.

70. Subramanian A, Tamayo P, Mootha VK, Mukherjee S, Ebert BL, Gillette MA, Paulovich A, Pomeroy SL, Golub TR, Lander ES et al.: Gene set enrichment analysis: a knowledge-based approach for interpreting genome-wide expression profiles. Proc Natl Acad Sci U S A 2005, 102(43):15545–15550.

71. Storey JD, Tibshirani R: Statistical significance for genomewide studies. Proc Natl Acad Sci U S A 2003, 100(16):9440–9445.

72. Wu Z, Irizarry RA, Gentleman R, Murillo FM, Spencer F: A Model Based Background Adjustment for Oligonucleotide Expression Arrays. J Amer Stat Assoc 2004, 99(468):909–917.

73. Benjamini Y, Hochberg Y: Controlling the false discovery rate: a practical and powerful approach to multiple testing. J Roy Statist Soc Ser B 1995, 57:289--300.

74. Wall DP, Pivovarov R, Tong M, et al. Genotator: a disease-agnostic tool for genetic annotation of disease. BMC medical genomics 2010; 3(1): 50.

75. Wall DP, Kudtarkar P, Fusaro VA, Pivovarov R, Patil P, Tonellato PJ. Cloud computing for comparative genomics. BMC bioinformatics 2010; 11(1): 259.

76. Wall D, Esteban F, Deluca T, et al. Comparative analysis of neurological disorders focuses genome-wide search for autism genes. Genomics 2009; 93(2): 120–9.

77. Varma M, Paskov KM, Jung J-Y, et al. Outgroup Machine Learning Approach Identifies Single Nucleotide Variants in Noncoding DNA Associated with Autism Spectrum Disorder. PSB; 2019; 2019. p. 260–71.

78. Vardarajan B, Eran A, Jung J, Kunkel L, Wall D. Haplotype structure enables prioritization of common markers and candidate genes in autism spectrum disorder. Translational psychiatry 2013; 3(5): e262–e.

79. Ruzzo EK, Pérez-Cano L, Jung J-Y, et al. Inherited and de novo genetic risk for autism impacts shared networks. Cell 2019; 178(4): 850–66. e26.

80. Nelson TH, Jung J-Y, DeLuca TF, Hinebaugh BK, St Gabriel KC, Wall DP. Autworks: a cross-disease network biology application for Autism and related disorders. BMC medical genomics 2012; 5(1): 56.

81. Li J, Ma Z, Shi M, et al. Identification of human neuronal protein complexes reveals biochemical activities and convergent mechanisms of action in autism spectrum disorders. Cell systems 2015; 1(5): 361–74.

82. Jung J, Kohane IS, Wall D. Identification of autoimmune gene signatures in autism. Translational psychiatry 2011; 1(12): e63–e.

83. Herbeck JT, Wall DP. Converging on a general model of protein evolution. TRENDS in Biotechnology 2005; 23(10): 485–7.

84. Goldfeder RL, Wall DP, Khoury MJ, Ioannidis JP, Ashley EA. Human genome sequencing at the population scale: a primer on high-throughput DNA sequencing and analysis. American journal of epidemiology 2017; 186(8): 1000–9.

85. Fusaro VA, Patil P, Gafni E, Wall DP, Tonellato PJ. Biomedical cloud computing with amazon web services. PLoS computational biology 2011; 7(8): e1002147.

86. Esteban FJ, Wall DP. Using game theory to detect genes involved in Autism Spectrum Disorder. Top 2011; 19(1): 121–9.

87. Diaz-Beltran L, Esteban FJ, Varma M, Ortuzk A, David M, Wall DP. Cross-disorder comparative analysis of comorbid conditions reveals novel autism candidate genes. BMC genomics 2017; 18(1): 315.

88. Diaz-Beltran L, Esteban F, Wall D. A common molecular signature in ASD gene expression: following Root 66 to autism. Translational psychiatry 2016; 6(1): e705–e.

89. Diaz-Beltran L, Cano C, Wall DP, Esteban FJ. Systems biology as a comparative approach to understand complex gene expression in neurological diseases. Behavioral Sciences 2013; 3(2): 253–72.

90. David MM, Enard D, Ozturk A, et al. Comorbid analysis of genes associated with autism spectrum disorders reveals differential evolutionary constraints. PloS one 2016; 11(7): e0157937.

91. David MM, Babineau BA, Wall DP. Can we accelerate autism discoveries through crowdsourcing? Research in Autism Spectrum Disorders 2016; 32: 80–3.

92. Cano C, Monaghan T, Blanco A, Wall DP, Peshkin L. Collaborative text-annotation resource for disease-centered relation extraction from biomedical text. Journal of biomedical informatics 2009; 42(5): 967–77.

93. Altman RB, Prabhu S, Sidow A, et al. A research roadmap for next-generation sequencing informatics. Science translational medicine 2016; 8(335): 335ps10–ps10.

94. Abbas, H., et al. Machine learning for early detection of autism (and other conditions) using a parental questionnaire and home video screening. in 2017 IEEE International Conference on Big Data (Big Data). 2017. IEEE.

95. Abbas, H., et al., Machine learning approach for early detection of autism by combining questionnaire and home video screening. Journal of the American Medical Informatics Association, 2018. 25(8): p. 1000–1007.

96. Abbas, H., et al., Multi-modular Ai Approach to Streamline Autism Diagnosis in Young children. Scientific reports, 2020. 10(1): p. 1–8.

97. Albert, N., et al., GapMap: enabling comprehensive autism resource epidemiology. JMIR public health and surveillance, 2017. 3(2): p. e27.

98. Altman, R.B., et al., A research roadmap for next-generation sequencing informatics. Science translational medicine, 2016. 8(335): p. 335ps10–335ps10.

99. Cano, C., et al., Collaborative text-annotation resource for disease-centered relation extraction from biomedical text. Journal of biomedical informatics, 2009. 42(5): p. 967–977.

100. Chrisman, B., et al. Analysis of Sex and Recurrence Ratios in Simplex and Multiplex Autism Spectrum Disorder Implicates Sex-Specific Alleles as Inheritance Mechanism. in 2018 IEEE International Conference on Bioinformatics and Biomedicine (BIBM). 2018. IEEE.

101. Cui, J., et al., Phylogenetically informed logic relationships improve detection of biological network organization. BMC bioinformatics, 2011. 12(1): p. 1–11.

102. Cui, J., et al., Detecting biological network organization and functional gene orthologs. Bioinformatics, 2011. 27(20): p. 2919–2920.

103. Daniels, J., et al., Feasibility testing of a wearable behavioral aid for social learning in children with autism. Applied clinical informatics, 2018. 9(1): p. 129.

104. Daniels, J., et al., Exploratory study examining the at-home feasibility of a wearable tool for social-affective learning in children with autism, npj Digital Medicine 1. 2018, Article.

105. Daniels, J., et al., Exploratory study examining the at-home feasibility of a wearable tool for social-affective learning in children with autism. NPJ digital medicine, 2018. 1(1): p. 1–10.

106. David, M.M., B.A. Babineau, and D.P. Wall, Can we accelerate autism discoveries through crowdsourcing? Research in Autism Spectrum Disorders, 2016. 32: p. 80–83.

107. David, M.M., et al., Comorbid analysis of genes associated with autism spectrum disorders reveals differential evolutionary constraints. PloS one, 2016. 11(7): p. e0157937.

108. Diaz-Beltran, L., et al., Systems biology as a comparative approach to understand complex gene expression in neurological diseases. Behavioral Sciences, 2013. 3(2): p. 253–272.

109. Diaz-Beltran, L., F. Esteban, and D. Wall, A common molecular signature in ASD gene expression: following Root 66 to autism. Translational psychiatry, 2016. 6(1): p. e705–e705.

110. Diaz-Beltran, L., et al., Cross-disorder comparative analysis of comorbid conditions reveals novel autism candidate genes. BMC genomics, 2017. 18(1): p. 315.

111. Duda, M., J. Daniels, and D.P. Wall, Clinical evaluation of a novel and mobile autism risk assessment. Journal of autism and developmental disorders, 2016. 46(6): p. 1953–1961.

112. Duda, M., et al., Crowdsourced validation of a machine-learning classification system for autism and ADHD. Translational psychiatry, 2017. 7(5): p. e1133–e1133.

113. Duda, M., J. Kosmicki, and D. Wall, Testing the accuracy of an observation-based classifier for rapid detection of autism risk. Translational psychiatry, 2015. 5(4): p. e556–e556.

114. Duda, M., T. Nelson, and D.P. Wall, Cross-pollination of research findings, although uncommon, may accelerate discovery of human disease genes. BMC Medical Genetics, 2012. 13(1): p. 114.

115. Duda, M., et al., Brain-specific functional relationship networks inform autism spectrum disorder gene prediction. Translational psychiatry, 2018. 8(1): p. 1–9.

116. El-Kalioby, M., et al. Personalized cloud-based bioinformatics services for research and education: use cases and the elasticHPC package. in BMC bioinformatics. 2012. BioMed Central.

117. Esteban, F.J. and D.P. Wall, Using game theory to detect genes involved in Autism Spectrum Disorder. Top, 2011. 19(1): p. 121–129.

118. Fusaro, V.A., et al., The potential of accelerating early detection of autism through content analysis of YouTube videos. PLOS one, 2014. 9(4): p. e93533.

119. Gafni, E., et al., COSMOS: Python library for massively parallel workflows. Bioinformatics, 2014. 30(20): p. 2956–2958.

120. Goldfeder, R.L., et al., Human genome sequencing at the population scale: a primer on high-throughput DNA sequencing and analysis. American journal of epidemiology, 2017. 186(8): p. 1000–1009.

121. Gupta, A., et al. Coalitional game theory as a promising approach to identify candidate autism genes. in PSB. 2018.

122. Haber, N., et al. A practical approach to real-time neutral feature subtraction for facial expression recognition. in 2016 IEEE Winter Conference on Applications of Computer Vision (WACV). 2016. IEEE.

123. Haber, N., C. Voss, and D. Wall, Making emotions transparent: Google Glass helps autistic kids understand facial expressions through augmented-reaiity therapy. IEEE Spectrum, 2020. 57(4): p. 46–52.

124. Herbeck, J.T. and D.P. Wall, Converging on a general model of protein evolution. TRENDS in Biotechnology, 2005. 23(10): p. 485–487.

125. Jung, J., I.S. Kohane, and D. Wall, Identification of autoimmune gene signatures in autism. Translational psychiatry, 2011. 1(12): p. e63–e63.

126. Jung, J.-Y., et al., A literature search tool for intelligent extraction of disease-associated genes. Journal of the American Medical Informatics Association, 2014. 21(3): p. 399–405.

127. Kalantarian, H., et al., A mobile game for automatic emotion-labeling of images. IEEE Transactions on Games, 2018.

128. Kim, I., et al., Cloud computing for comparative genomics with windows azure platform. Evolutionary Bioinformatics, 2012. 8: p. EBO. S9946.

129. Kline, A., et al., Superpower glass. GetMobile: Mobile Computing and Communications, 2019. 23(2): p. 35–38.

130. Levy, S., et al., Sparsifying machine learning models identify stable subsets of predictive features for behavioral detection of autism. Molecular autism, 2017. 8(1): p. 65.

131. Li, J., et al., Identification of human neuronal protein complexes reveals biochemical activities and convergent mechanisms of action in autism spectrum disorders. Cell systems, 2015. 1(5): p. 361–374.

132. Nag, A., et al., Toward Continuous Social Phenotyping: Analyzing Gaze Patterns in an Emotion Recognition Task for Children With Autism Through Wearable Smart Glasses. Journal of Medical Internet Research, 2020. 22(4): p. e13810.

133. Nelson, T.H., et al., Autworks: a cross-disease network biology application for Autism and related disorders. BMC medical genomics, 2012. 5(1): p. 56.

134. Paskov, K.M. and D.P. Wall, A low rank model for phenotype imputation in autism spectrum disorder. AMIA Summits on Translational Science Proceedings, 2018. 2018: p. 178.

135. Saxe, G.N., et al., A complex systems approach to causal discovery in psychiatry. PloS one, 2016. 11(3): p. e0151174.

136. Sivakumaran, T.A., et al., Conservation of the RB1 gene in human and primates. Human mutation, 2005. 25(4): p. 396–409.

137. Souilmi, Y., et al., Scalable and cost-effective NGS genotyping in the cloud. BMC medical genomics, 2015. 8(1): p. 1–9.

138. Sun, M.W., et al., Coalitional Game Theory Facilitates Identification of Non-Coding Variants Associated With Autism. Biomedical Informatics Insights, 2019. 11: p. 1178222619832859.

139. Sun, M.W., et al., Game theoretic centrality: a novel approach to prioritize disease candidate genes by combining biological networks with the Shapley value. BMC bioinformatics, 2020. 21(1): p. 1–10.

140. Vardarajan, B., et al., Haplotype structure enables prioritization of common markers and candidate genes in autism spectrum disorder. Translational psychiatry, 2013. 3(5): p. e262–e262.

141. Varma, M., et al. Outgroup Machine Learning Approach Identifies Single Nucleotide Variants in Noncoding DNA Associated with Autism Spectrum Disorder. in PSB. 2019.

142. Venkataraman, G.R., et al. De novo mutations in autism implicate the synaptic elimination network. in PACIFIC SYMPOSIUM ON BIOCOMPUTING 2017. 2017.

143. Voss, C., et al., Effect of wearable digital intervention for improving socialization in children with autism spectrum disorder: a randomized clinical trial. JAMA pediatrics, 2019. 173(5): p. 446–454.

144. Voss, C., et al. Superpower glass: delivering unobtrusive real-time social cues in wearable systems. in Proceedings of the 2016 ACM International Joint Conference on Pervasive and Ubiquitous Computing: Adjunct. 2016.

145. Wall, D.P., Use of the nuclear gene glyceraldehyde 3-phosphate dehydrogenase for phylogeny reconstruction of recently diverged lineages in Mitthyridium (Musci: Calymperaceae). Molecular phylogenetics and evolution, 2002. 25(1): p. 10–26.

146. Wall, D.P. and J.T. Herbeck, Evolutionary patterns of codon usage in the chloroplast gene rbcL. Journal of molecular evolution, 2003. 56(6): p. 673–688.

147. Wall, D.P. and P.J. Tonellato, The future of genomics in pathology. F1000 medicine reports, 2012. 4.

148. Wall, D.P. and P.J. Tonellato, Deriving clinical action from whole-genome analysis. Personalized Medicine, 2012. 9(3): p. 247–252.

149. Washington, P., et al., Precision Telemedicine through Crowdsourced Machine Learning: Testing Variability of Crowd Workers for Video-Based Autism Feature Recognition. Journal of personalized medicine, 2020. 10(3): p. 86.

150. Washington, P., et al. Feature selection and dimension reduction of social autism data. in Pac Symp Biocomput. 2020.

